# Structure guided significance testing correction for hydrogen deuterium exchange mass spectrometry

**DOI:** 10.1101/2025.06.16.659914

**Authors:** Oliver M. Crook

## Abstract

Hydrogen deuterium exchange mass spectrometry (HDX-MS) is a powerful technique to probe changes in protein structural dynamics. In differential settings, HDX-MS compares dynamics between protein states, such as conformational changes resulting from antibody-antigen binding or the effects of mutations. As the method becomes more high-throughput, the number of comparisons between peptides and states grows, creating a multiple hypothesis testing challenge where some observed changes may result from statistical randomness rather than biological differences. While this problem can be addressed by applying multiple hypothesis testing correction techniques like false discovery rate (FDR) control, current methods assume statistical independence - an assumption violated by peptide overlap and the influence of protein structure. Here, we develop a structural false discovery rate (sFDR) that accounts for these dependencies by integrating sequence and structural information to estimate the effective number of independent tests. Our approach significantly improves statistical power to detect genuine changes in protein dynamics measured by HDX-MS, as demonstrated through extensive validation using both simulated and experimental datasets. The sFDR method maintains robustness to structural uncertainty, making it applicable even when only predicted structures are available. This advancement enables more sensitive detection of conformational changes in challenging systems with subtle dynamic differences and reduces the number of replicates needed to obtain confidence in results. The method is easily accessible through a Google Collab Notebook and can be completed in minutes.

## Introduction

Understanding protein conformational dynamics and interaction-induced structural perturbations is essential for characterising protein function. Hydrogen-deuterium exchange mass-spectrometry (HDX-MS) provides a powerful approach to investigate these dynamics by measuring the rate at which amide hydrogens exchange with deuterium in solution (Orengo *et al*., 1999; Engen, 2009; Chalmers *et al*., 2011; Houde *et al*., 2011; Masson *et al*., 2017; Sauve *et al*., 2018). The exchange rate is influenced by structural elements including secondary and higher-order arrangements, as well as solvent accessibility (Katta and Chait, 1993; Konermann *et al*., 2011; Kammari and Topp, 2020; Jia *et al*., 2020). This methodology is grounded in the Linderstrøm-Lang theory of local folding and unfolding events (James *et al*., 2021). HDX-MS applications are numerous: mapping binding interfaces in antibody-antigen complexes (Tsai *et al*., 2020); investigating structural stabilization through small molecule interactions and their allosteric consequences (Kielkopf *et al*., 2019); assessing the structural impact of amino acid mutations in transporters (Jia *et al*., 2020); and contributing to rational vaccine design (Gorman *et al*., 2016).

Differential HDX-MS compares deuterium incorporation patterns between multiple protein states using matched peptides across samples. The mass difference between deuterium and hydrogen enables accurate measurements using mass spectrometry. When identifying antibody epitopes, for example, researchers typically observe decreased HDX rates in peptides at binding interfaces (Tsai *et al*., 2020). However, changes in HDX rates may result from both biological perturbations and measurement variability, which requires statistical assessment of peptide-level significance. Statistical approaches include global thresholds, mixed effects models (Hourdel *et al*., 2016), hybrid tests (Hageman and Weis, 2019), moderated statistics (Claesen *et al*., 2021), functional data analysis (Crook *et al*., 2022), and Bayesian methods (Crook *et al*., 2023, 2024). Most of these methodologies subsequently apply False Discovery Rate (FDR) control to adjust p-values to q-values, accounting for multiple testing (Benjamini and Hochberg, 1995). This adjustment, however, assumes test independence which is an invalid assumption given the inherent sequence overlap between peptides and the influence of the three-dimensional protein structure on exchange dynamics. These statistical limitations can significantly impact biological interpretations, particularly in regions with subtle structural dynamics.

To address these statistical limitations, we introduce the structural False Discovery Rate

(sFDR), a novel approach that controls for multiple hypothesis testing while accounting for the inherent spatial correlations in HDX-MS data. Unlike conventional methods, sFDR integrates peptide coordinates with protein structural information, allowing experimentally determined or computationally predicted structures, to calculate an effective number of independent tests *M*_eff_. This framework accommodates structural uncertainty through confidence metrics such as predicted local distance difference test (pLDDT) scores, providing robust statistical control even when using predicted models (Jumper *et al*., 2021).

Through comprehensive simulation studies, we demonstrate that sFDR maintains strict control of false positive rates even under scenarios of substantial structural noise, confirming the reliability of our approach with predicted protein structures. Application to experimental datasets reveals significantly improved detection of biologically relevant conformational changes compared to traditional FDR approaches. Notably, these improvements are most pronounced in regions with complex structural arrangements where conventional methods often fail. The approach is made available using a simple to use Google Collab Notebook: https://colab.research.google.com/drive/1TB_UhK4cy6ogJfH-tt1pvNkGw49ekjWT?usp=sharing and Github: https://github.com/ococrook/hdx-sFDR

## Results

### Structure guided significance testing

We present a statistical framework that models the inherent correlations in HDX-MS data arising from both sequence overlap and structural proximity (Fig.1A). Our approach constructs a weight matrix that integrates three key components: one, sequence relationships quantified as mid-point distances between peptides scaled by their fractional overlap; two, structural proximity measured as the Euclidean distance between peptide centroids in three-dimensional space; and three structural confidence represented by the mean per-residue confidence values for each peptide. These components are combined through a weighted average controlled by a mixing parameter *α* (Fig.1B). To determine the effective number of independent tests *M*_eff_, we normalise this weight matrix to a correlation matrix and perform eigenvalue decomposition (Fig.1C). By summing the capped eigenvalues (maximum value of 1), we obtain *M*_eff_, which is typically substantially smaller than the nominal number of tests, thereby increasing statistical power. This approach enables the generation of structure-aware q-values that maintain appropriate type I error control (false positives) while accounting for the complex dependencies inherent in HDX-MS data (Fig.1D). Detailed mathematical formulations are provided in the Methods section.

**Figure 1:**
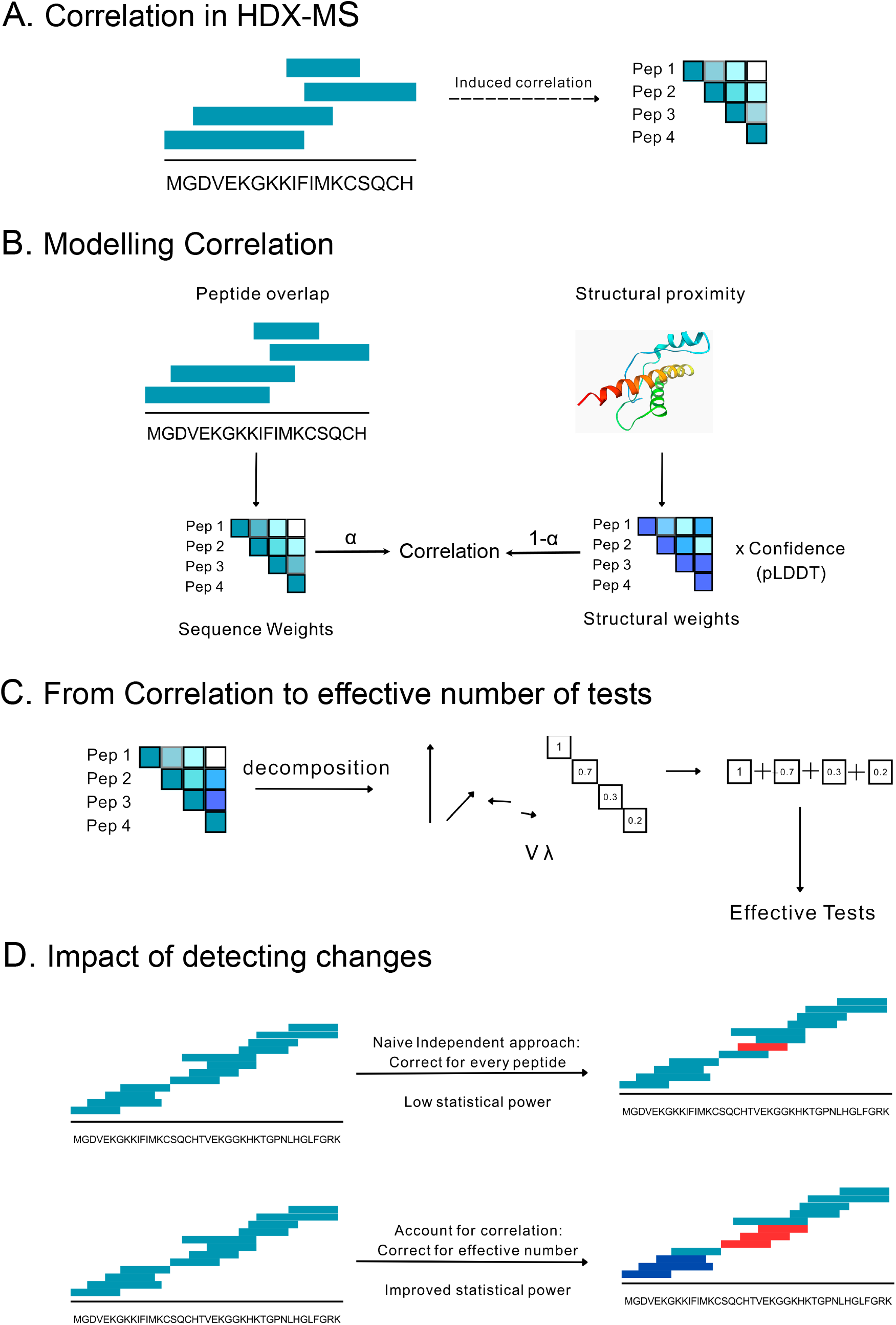
Overview of structural False Discovery Rate (sFDR) methodology. A) Correlation in HDX-MS data arises from peptide sequence overlap and structural proximity. Overlapping peptides (shown as horizontal bars) covering the amino acid sequence MGDVEKGKKIFIMKCSQCH exhibit induced correlations due to shared residues, represented in the correlation matrix heatmap where warmer colours indicate stronger correlations between peptide pairs. (B) Modelling correlation through weighted integration of sequence and structural information. Sequence weights are calculated from peptide overlap and mid-point distances, while structural weights are derived from 3D Euclidean distances between peptide centroids in the protein structure. These components are combined using mixing parameter *α*, with structural weights further modulated by confidence scores (pLDDT values from structure prediction). The final correlation matrix integrates both dimensions: Correlation ≈ *α* × (Sequence Weights) + (1-*α*) × (Structural Weights × Confidence).(C) Calculation of effective number of tests from correlation structure. Eigenvalue decomposition (*V λ*) of the correlation matrix reveals that many eigenvalues are substantially less than 1, indicating dependencies between tests. The effective number of tests (*M*_eff_) is calculated by summing eigenvalues capped at 1, typically yielding *M*_eff_ *<< M* (total peptides), thereby reducing statistical stringency. (D) Impact on detecting clustered biological effects. Traditional FDR correction assumes independence between all peptides, leading to overly conservative corrections and low statistical power to detect spatially clustered effects (top panel). The sFDR approach accounts for correlations, improving power to detect biologically meaningful clustered changes while maintaining appropriate false positive control (bottom panel). Red and dark blue peptides indicate significant differences detected by each method.

### Structural FDR reduces number of tests and is robust to structural uncertainty

To demonstrate the practical impact of sFDR, we analysed Cytochrome C as a model system. Using a peptide map generated from a Nepenthesin II digestion (Crook *et al*., 2024) (yielding 74 overlapping peptides) and five structural models predicted by AlphaFold 3 (Abramson *et al*., 2024), we computed the required correlation matrices. The resultant matrix (Fig. 2 A) reveals distinct correlation clusters that correspond to structural domains and sequence overlap within the protein. Eigenvalue decomposition of this matrix, Fig. 2B, shows that many eigenvalues are substantially less than one, indicating that the conventional approach of using the total peptide count as the number of tests would significantly overestimate statistical stringency. When applied to the top-ranked AlphaFold model, our method calculated an effective number of tests *M*_eff_ of approximately 16.0, representing a 4.6-fold reduction from the nominal count. Furthermore, *M*_eff_ values derived from all five AlphaFold models were remarkably consistent (closest integer was always 16), demonstrating that our approach is robust to structural prediction uncertainty and suggesting that even lower-ranked structural models provide sufficient information for reliable statistical correction.

**Figure 2:**
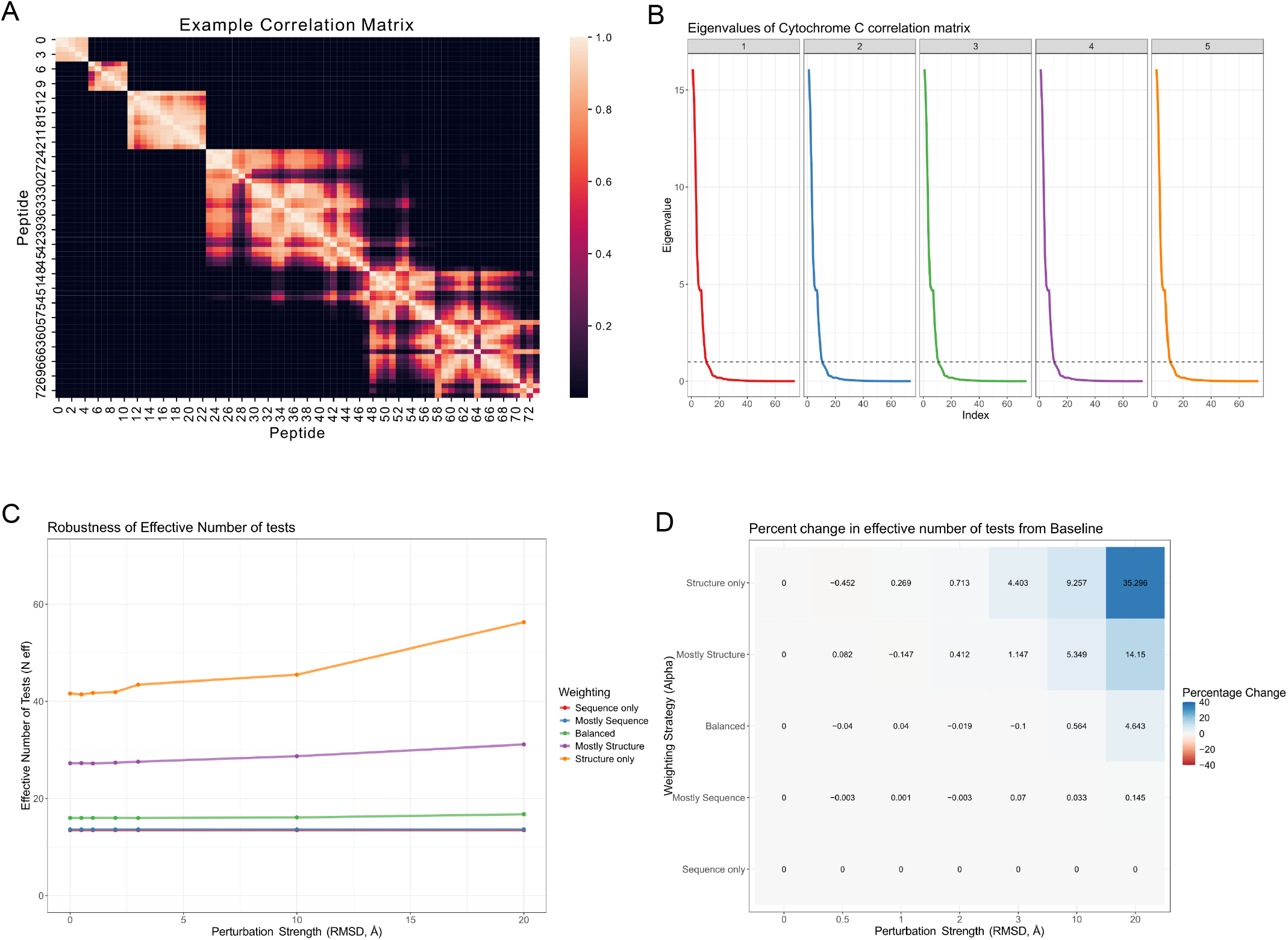
Properties of sFDR. (A) An example correlation matrix derived from our method. There is a clear clustering structure in the correlation matrix. (B) The eigen-value spectrum of the correlation matrix derived from Cytochrome C data. The dashed line indicates a large number of eigenvalues are less than 1 and hence the number of tests is over estimated. (C) A line graph showing how the effective number of tests is a function of weighting between sequence and structural information. Using more structure information results in a higher number of effective tests. The number of effective also increases as a function of perturbation strength to the initial structure. The more structural information used the greater the impact of structural perturbation. (D) A heatmap quantifying the percentage change in the number of effective tests under different scenarios (weighting and structural noise).

To evaluate sFDR’s sensitivity to structural uncertainty, we systematically assessed its performance under increasing levels of structural perturbation (measured as RMSD in Å) across the full spectrum of the mixing parameter *α*, using the same Cytochrome C system. We tested five *α* values ranging from 0 (structure-only weighting) to 1 (sequence-only weighting), including intermediate values of 0.2, 0.5 and 0.8. As anticipated, structural noise affected *M*_eff_ only when structural information was incorporated into the weighting scheme (*α <* 1). The structure-only approach (*α* = 0) exhibited high sensitivity to structural perturbations, with *M*_eff_ increasing to approximately 60 under severe distortion (20Å RMSD). In contrast, the balanced weighting approach (*α* = 0.5) demonstrated remarkable resilience, showing only minor sensitivity (≤ 5% change in *M*_eff_) even under substantial structural perturbation. This finding suggests that balanced incorporation of both sequence and structural information provides optimal robustness to structural uncertainty, supporting the reliable application of sFDR with predicted protein structures.

### Structural FDR improves sensitivity and reduces false positives

We next evaluated sFDR’s ability to enhance statistical sensitivity while maintaining strict false positive control. Using the Cytochrome C system, we designed simulation experiments with both null and alternative hypotheses. For the null scenario (no differential HDX), we sampled p-values from a uniform distribution across all peptides. For the alternative hypothesis, we designated a contiguous region (e.g. residues 25-30) as differentially exchanged, generating p-values from U(0, 0.001) for peptides overlapping this region. We then compared multiple testing correction using our structure-aware approach (weighted) versus the conventional Benjamini-Hochberg (original) procedure (Benjamini and Hochberg, 1995).

At a stringent q-value threshold of 0.001, sFDR substantially outperformed the conventional approach in sensitivity (Fig. 3A and B). By systematically varying the mixing parameter *α*, we demonstrated that a balanced integration of structural and sequence information (*α* = 0.5) achieved optimal performance. The conventional method failed to detect any true differences at this threshold, unable to leverage the spatial clustering of effects (Fig. 3B). Importantly, in a negative control scenario with no simulated differences, sFDR maintained appropriate type I error control without generating spurious positives (Fig. 3C). Further analysis of FDR control revealed critical differences between the approaches.

**Figure 3:**
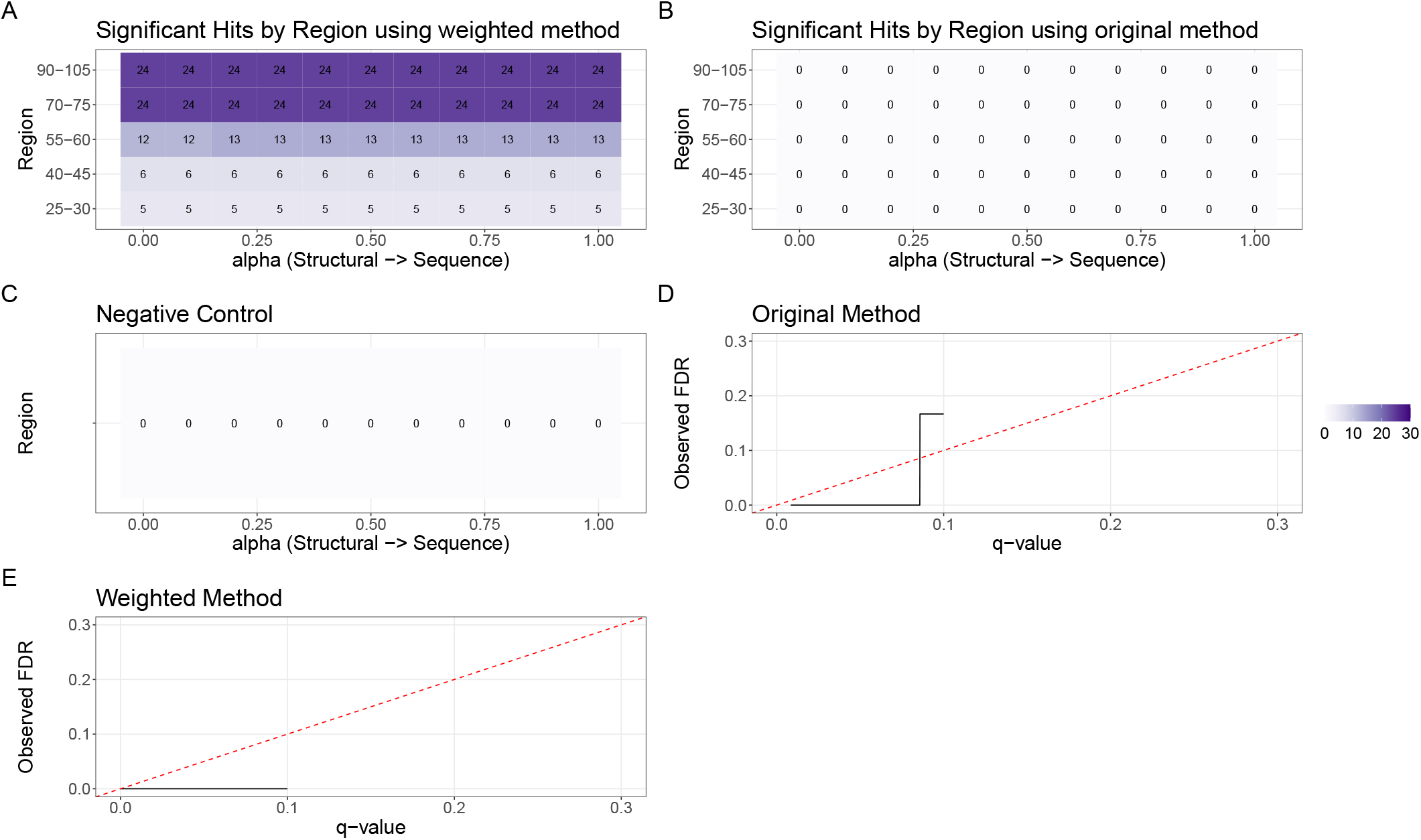
Simulation study using Cytochrome C. (A) Number of peptides that were declared signifiant for different regions selected to have structural changes as a function of the amount of structural information used in the weighting approach. (B) As for A but assuming all peptides are independent. (C) Negative control using the weighted method to calculate q-values but no peptides were simulated to be significantly different. No peptides should be declared significant. (D) Standard q-values against observed FDR. Desirable q-values conservatively estimated the FDR and hence should be below the dashed red line. (E) As for D but using the weighted q-values.

The conventional method occasionally generated false positives when statistically random small p-values occurred in isolation (Fig. 3D). In contrast, sFDR effectively exploited the expected spatial clustering of biological effects, producing appropriately conservative q-values that better reflected the underlying structural relationships (Fig. 3E). This demonstrates that sFDR not only improves sensitivity but also down-weights isolated and spurious effects in HDX-MS experiments.

### Epitope mapping HOIP dAbs using HDX-MS

Epitope mapping in standard application of HDX-MS, whereby binding epitopes are identified by protection signatures because the antibody stabilises the hydrogen bond network at the binding site. HOIP, an E3 ubiquitin ligase, is a pharmaceutical target due it’s role in immune signalling (Kirisako *et al*., 2006). Previous authors have performed HDX-MS experiments for a ring-between-ring construct of HOIP, hence HOIP-RBR, in complex with single domain antibodies and in unbound state (Tsai *et al*., 2020). A standard HDX-MS workflow was performed with MS performed using a Water’s Synapt G2-Si instrument. HDX-MS measurements were taken at 0, 30 and 300s post-exposure to heavy water, for a number of dAbs at different molar concentrations (Tsai *et al*., 2020).

In this experiment replicates were sacrificed in order to profile a greater number of antibodies. However, this means in several cases the epitope and potentially other dynamic effects were left unidentified due to loss of statistical power. Even with methods that improve power by exploiting the time dimension of the data, the epitopes are still unidentified in some cases (Crook *et al*., 2022). Here, we focus on three examples, dAb13, dAb25 and dAb6, as they cover a range of scenarios.

To analyse these experiments the statistical method ‘hdxstats’ was used as described in previous work (Crook *et al*., 2022). Subsequently, any *p*-values from this method were corrected using sFDR exploiting peptide overlap, the highest ranked Alphafold 3 structure of HOIP-RBR and the corresponding pLDDT (Abramson *et al*., 2024). The results are visualised in figure Fig. 4 with the peptides identified as significant by ‘hdxstats’ (FDR < 0.05) denoted by a dark green star whilst those peptides with sFDR < 0.05 are marked by a black star. Noting the only difference between the analysis is the use of FDR or sFDR with the underlying statistical testing remaining identical.

**Figure 4:**
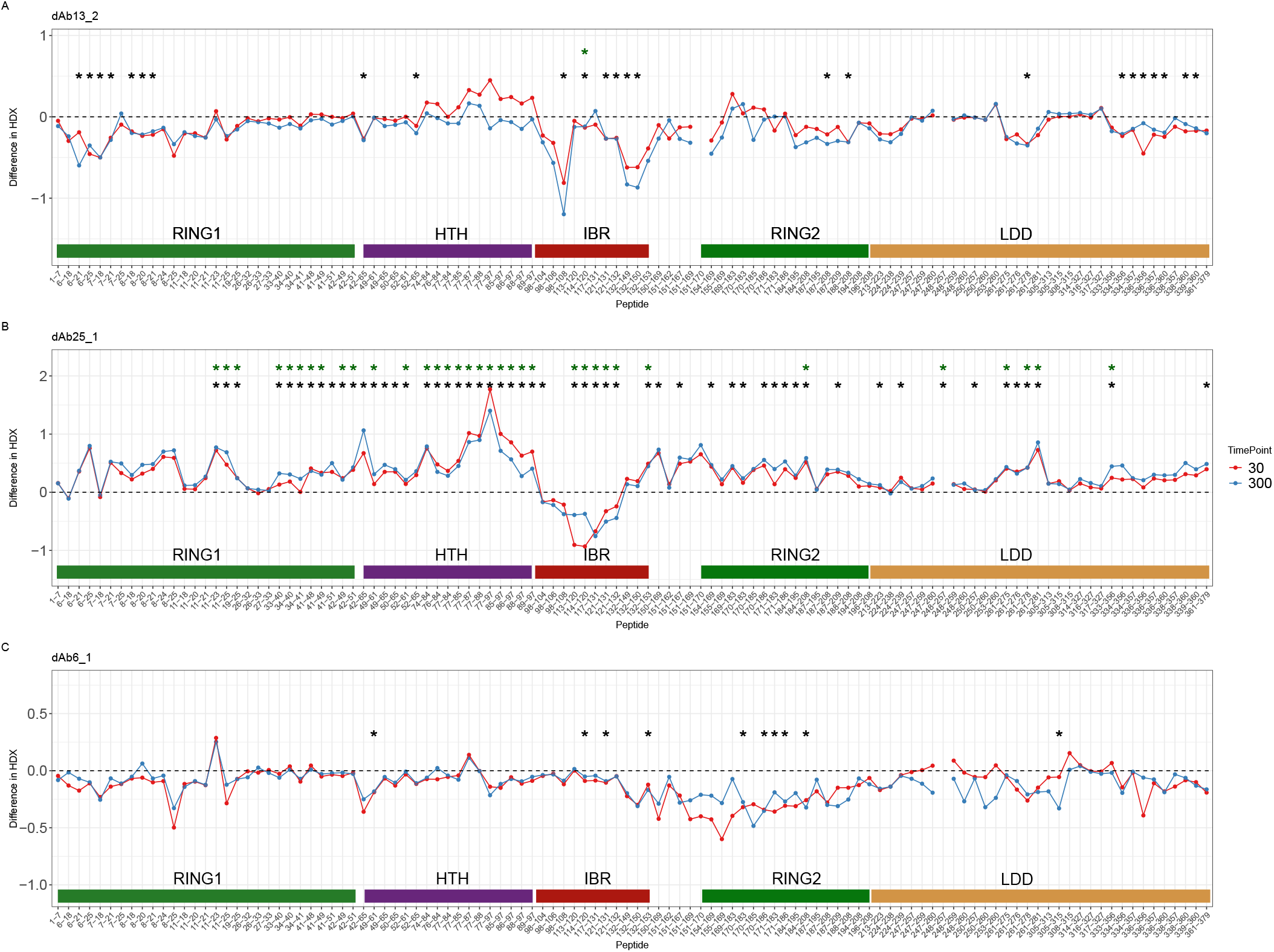
Epitope mapping HOIP-RBR. (A) A butterfly plot with peptides mapped against pointwise deuterium uptake differences in units of Daltons. Structural regions are highlighted using a colour bar. The dark green stars indicate peptides that are significant using standard FDR procedures whilst the black stars indicate peptides that are significant using the sFDR approach. The difference in HDX is between unbound HOIP-RBR and HOIP bound to dAb13 at 2 molar concentration. (B) As for A but dAb25 at 1 molar concentration is used (C) As for A but dAb6 at 1 molar concentration is used.

Fig. 4A shows a dramatic scenario were previous approaches only identified a single significant peptide, whereas using our sFDR identifies 25 peptides which cluster in the sequence dimension as expected. Where previous analysis would have been unable to identify a putative epitope, here we have evidence in support of a non-linear epitope on the right and left-sides of the in-between ring (IBR). Distal regions of protection also allow us to identify previously uncharacterised protein dynamics as consequence of binding. As observed with other dAbs, binding at the IBR appears to induce a closed or compact conformation as evidenced by a reduction in HDX in the RING1 and linear-determining domain (LDD) regions.

Fig. 4B shows an example were previous analysis was able to identify a continuous epitope in the central IBR region and induces a open conformation of HOIP-RBR as evidenced by the increase in HDX in the bound state. Here, employing sFDR provides further confidence to this observation and gives statistical support to the de-protection observed in RING2, suggestive of a global open conformation rather than local structural rearrangements in the HTH that could increase HDX.

In the case of dAb6, Fig.4C, prior approaches were unable to identify an epitope despite dAb6 being one of only two antibodies in the previous study to be inhibitory of both UbcH7 (strictly cysteine reactive E2-conjugating enzyme) and UbcH5C (lysine reactive E2-conjugating enzyme) activity (Tsai *et al*., 2020). Here, we provide statistical support for the dAb6 epitope to be shifted to right of the IBR in the RING2 region. dAb6’s inhibitory ability suggests that it is able to block the catalytic cysteine at residue 188 (855 on full protein) and the ubiquitin donor site in the IBR. This may suggest that dAb6 binding might trap HOIP-RBR in a closed conformation or encourage structural rearrangements that cause the catalytic cysteine to become occluded.

These results highlight the benefit of sFDR to uncover epitopes and structural dynamics in HDX-MS data were previous methods have failed to identify the epitope or overlooked some structural dynamics. We include an additional seven epitope mapping experiments in the supplementary material to show broad applicability.

### HDX-MS of a structural variant experiment

A previous study used a structural variant model to define the detection limits of HDX-MS. Those authors performed HDX-MS on Maltose-binding protein (MBP) in wild-type (WT) and on a mutant protein, MBP-W169G, where the mutation site is buried in one of the lobes of MBP. The benefit of this system is that the mutant only subtly destabilises the structure and in this case the mutant slightly increases the hydrodynamic radius suggesting a more flexible and conformationally variable protein.

In their experiment the mutant was spiked into a sample containing WT protein generating a mixture distribution. To test our methodology in a challenging scenario, we looked for differences between a sample where 20% of the sample was mutant protein compared to a sample where 10% of the sample was mutant protein. The higher proportion of mutant suggests we should observe increases in HDX but the effects maybe subtle. To demonstrate our method is broadly applicable, we used a standard t-test to test whether the average difference between peptides measured in both experiments was different from zero, based on triplicate measurements of the system. To compare, the p-values where then adjusted for multiple testing to weighted q-values using our sFDR approach or to standard q-values using the Benjamini-Hochberg procedure. The highest rank Alphafold 3 predicted structure was used to derive the structural correlation.

The consequence of the differences between these approaches is visualised in Fig. 5A and Fig. 5B. Using standard q-values, we see a number of differences with many, as expected, in the lobe containing the structural variant. However, we also see some seemingly isolated differences especially at the earlier time points. For example, differences are observed at 30 seconds post exposure to heavy water in distal regions from the mutant where no overlapping peptides also have significant differences, suggestive of a false positive, (Fig. 5A and Fig. 5C). However, using the weighted q-values we proposed, we observe that these peptides are sometimes reinforced by overlapping peptides reducing the likelihood of a false positive. In addition, some peptides that were not significant previously are significant when the weighted q-values are used. These peptides frequently overlap with other significant peptides, suggesting that the weighted q-values effectively borrows statistical power from the spatial dimensions of the data. This approach adds statistical confidence to subtle effects that might otherwise be missed using standard methods. This effect is observed at all time-points with isolated effects down-weighted and clustered effects reinforced (see Fig. 5B and Fig. 5D)

**Figure 5:**
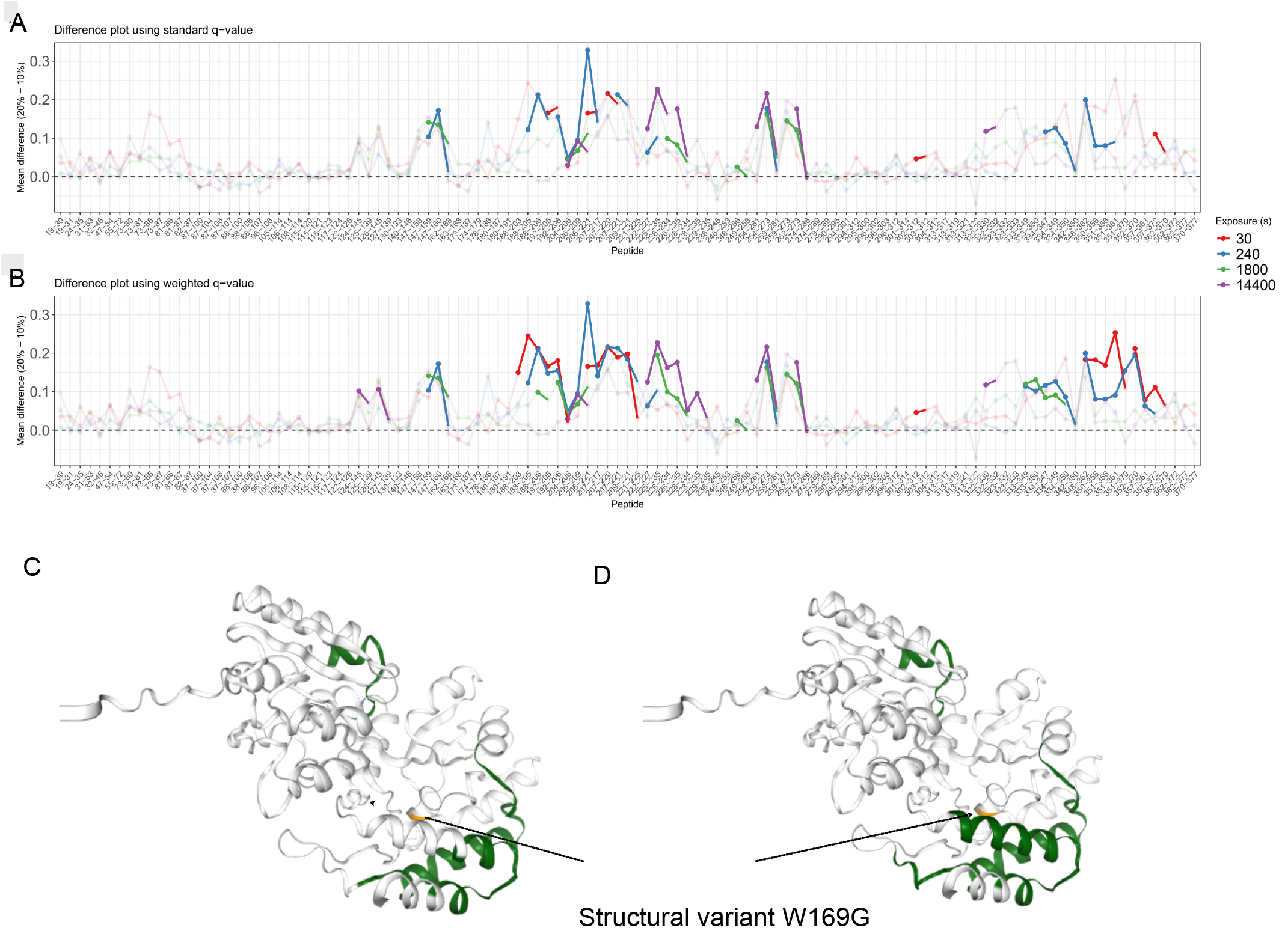
Structural variant differences using HDX-MS. (A) A masked butterfly plot with deuterium uptake differences between the 20% and 10% structural spike-in conditions. Using the standard q-values, we masked points that are not significant by making them transparent. The different colours indicate the measurement time points. (B) As for A but using the weighted q-values to assess significance. (C) Alphafold 3 predicted structure of MBP with peptides declared significant in A at 30 seconds exposure to heavy water coloured in dark green. The location of the structural variant is marked in orange. (D) As for C but the weighted q-values are used instead.

These findings demonstrate that our sFDR approach provides more biologically relevant results by reducing spurious signals while enhancing the detection of spatially consistent conformational changes, particularly for subtle structural variations.

## Discussion

We have presented a method to account for the non-independence of statistical tests performed in HDX-MS data analysis. Our method incorporates and accounts for the spatial components of HDX-MS both along the sequence axis and due to the 3 dimensional protein structure. The approach works by constructing a correlation matrix from the peptide overlap and structural proximity. The matrix can then be used to calculate the effective number of tests accounting for dependence using an eigen-decomposition or used as part of weighting schema.

Using empirical analysis across protein systems, we demonstrate the improvement in power and consistency this approach brings. Our simulations demonstrate control of the false discovery rate even when a predicted structure is used and there is significant structural noise. Indeed, structural noise will underestimate correlations returning our method to the more conservative scenario of assuming fully independent tests.

Two case-studies highlight the benefits of our approach. In an epitope mapping experiment we were able to identify previous unknown epitopes as well as structural dynamics. Even in cases were the epitope was clear from prior analysis our approach was able to give statistical confidence to dynamic that would have been overlooked otherwise. In a structural variant experiment, we find improved detection of subtle and consistent effects when using our methods.

While sFDR shows robust performance across diverse systems, several limitations are worth mentioning. For proteins with very sparse peptide coverage or very poor structural prediction could mean computed correlations are not meaningful. If the data are uncorrelated our approach gracefully degrades to standard FDR whilst users may need to perform sequence only analysis if the structural quality is particularly poor (*α* = 1). Since the main computational bottleneck is eigenvalue decomposition the approach scales to datasets with 1000’s of peptide, however will naturally become more computational intensive as size of datasets increases.

The correction methods we propose here are applicable to any prior statistical testing approach used and so is a highly general method. Future work could exploit correlations derived from molecular dynamics simulations. The methods maybe applicable in other proteomics methods that need to account for correlated tests.

## Methods

### Methodological Summary

We assume that the primary goal of statistical testing for HDX-MS is testing the null hypothesis that their is no difference in mean deuterium uptake for a given peptide between two or more conditions, for a given time point. This test would then be repeated across time points, for a total of *T*, and for each peptide in turn, for a total of *M*, typically results in *M* × *T* p-values that are subsequently adjusted to q-values using the Benjamini-Hochberg procedure to avoid inflation of the number of false positives. For example, without correcting p-values then *a* × 100% of p-values are smaller than *a* due to statistical sampling alone in the presence of no true biological effect. More complex methods to obtain a p-value are also valid for what follows including if they are derived from interaction effects of mixed effects models or from testing difference in kinetics using time dependent models.

However, unless the statistical approach explicitly models spatial correlations between peptides the standard approaches for correcting p-values will be overly conservative (reduce statistical power). First, consider two overlapping peptides the information shared between these peptides is a function of the extent of their overlap. They share information because they are measuring a shared subset of residues on which the exchange process is happening. Thus, a test for each peptide separately overlooks the fact that rejecting the null hypothesis for one peptide would increase our probability of rejecting the null hypothesis for the overlapping peptide. Two options are available to adjust for this effect. The first is corrected p-values which are obtained from computing the effective number tests using the correlation we obtain from using the structure and sequence overlap. Alternatively, we can compute weighted q-values by computing individual weights for p-values based on the correlation structure.

### Preliminaries

We first recall the false discovery rate (FDR) and the classical BH procedure (Benjamini and Hochberg, 1995). Suppose, we are testing a total of M hypothesis *H*_*i*_, *i* = 1, …, *M*. We assume that there are *m*_0_ null hypotheses with the remainder *m*_1_ = *M* − *m*_0_ where the alternative hypothesis is preferred. Let *V* be the number of false discoveries (falsely rejected) and R be the total number of rejected hypotheses. The FDR is given by

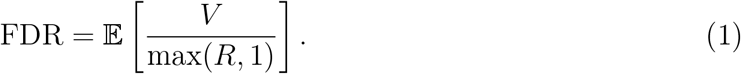

The BH procedure involves ordering the corresponding p-values *p*_(1)_, …, *p*_(*M*)_. Given a significance level *q* ∈ (0, 1), the BH procedure rejects all hypotheses (e.g. the alternative is preferred) for *p*_(*i*)_ ≤ *p*_(*k*)_, where

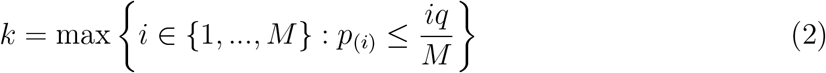

The original proof shows that the BH procedure controls the FDR at level *π*_0_*q* where *π*_0_ = *m*_0_*/M* is the proportion of true null hypothesis. Hence, given that *π*_0_ ≤ 1, the BH procedure actually controls the FDR more stringently than *q*.

### Corrected q-values

Following the methods of Li and Ji for genomics (Li and Ji, 2005), we assume we have access to a correlation matrix Σ for the peptides observed in HDX-MS analysis. The Σ represents that peptides are not in fact independent but rather correlated due to their peptide overlap and structural proximity. For narrative clarity, we assume we have access to this matrix and discuss calculating it in a later section.

As Li and Ji (2005) describe the idea is to calculate *M*_eff_, the effective number of tests from the correlation matrix. In the case of complete independence then the eigenvalues of the correlation matrix are all 1 and of course *M*_eff_ = *M*. Correlations will results in eigenvalues *λ*_*i*_ ∈ (0, 1). If *c* of the tests are identical then there is a *j* for which *λ*_*j*_ = *c*. Hence, we can decompose the spectrum, the set of eigenvalues, of the matrix Σ into their integral and non-integral parts. The integral parts count identical tests, which should count as 1 because an identical test has already been performed. The non-integral part represents a partially correlated test that should be counted fractionally between 0 and 1. Thus, the (Li and Ji, 2005) estimate of *M*_eff_ is

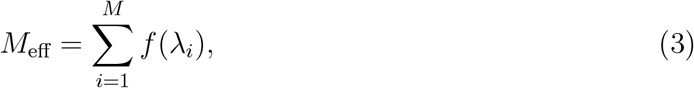

where *f* (*x*) = I(*x* ≥ 1) + (*x* − ⌊*x*⌋). I(*x* ≥ 1) is the indicator function giving 1 when *x* ≥ 1 and gives 0 otherwise, and ⌊*x*⌋ is the floor function, which returns the largest integer less than equal to *x. M*_eff_ then replaces *M* in the original BH procedure. Given that *M*_eff_ ≤ *M* power will be improved by accounting for these correlations. We refer to this as corrected q-values throughout the text.

### Weighted q-values

Alternatively, the correlation matrix can be used to adjust the p-values more directly. We wish to exploit that there may be regions of the protein for which nulls or alternatives are more likely than for other regions, without knowing this information in advance. If we could estimate *π*_0_, the proportion of nulls using 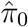 then the BH procedure can be applied to weighted p-values 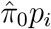. Methods that exploit using the data to improve power in this way are referred to *adaptive procedures*.

Following (Hu *et al*., 2010), we apply their adaptive two-stage (TST) group BH procedure, which we refer to as Adaptive TST GBH procedure. The rationale for this is that we can define groups in the data where we are more likely to reject the null hypothesis on certain groups than others. For example, peptides that are in an epitope. On each group, we will then obtain weights in the adaptive manner just described. We will explain later how to obtain peptide groups *g* = 1, …, *G* from the correlation matrix Σ. The procedure works as follows:

1. For the *p*-values in each group peptide group *g*, apply the standard BH method at level 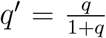, typically *q* = 0.05. Let *r*_*g*,1_ be the number of rejections and *n*_*g*_ the number of observations in that group.
2. For each group, compute the the TST estimator of *π*_*g*,0_:

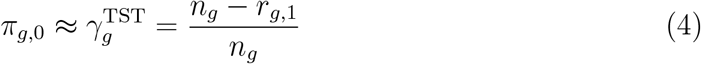
3. For each p-value in group *g*, calculate the weighted p-value 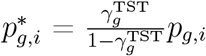. Capping *p*-values at 1.
4. Pool the p-values together and apply the standard BH procedure at level *q*^*′*^. Resulting in weighted q-values.

In practice, to avoid 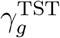 becoming too large or small, we replace 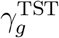 with 0.5 if the value is smaller than that and if 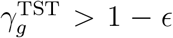 for *ϵ* = 10^*−*4^ then we replace the TST estimator with 1 − *ϵ* to avoid division by numbers very close to zero. We refer to the output of this approach as weighted q-values throughout the text.

To obtain groups from the correlation matrix Σ, we apply *K*-means clustering using the silhouette score to obtain the value of *K*.

### Computing the correlation matrix

In both scenarios of corrected and weighted the q-values require the use of the correlation/weight matrix that is constructed from the sequence and structure information. The correlation matrix construction follows a three-step logic (1) sequence overlap captures direct measurement dependencies (2) structural proximity captures biophysical dependences in HDX rates, and (3) confidence weighting ensures robust performance with predicted structures. These are combined and normalised to create a proper correlation matrix. More mathematically, the weight matrix is obtained by computing sequence weights *W*_seq_, structure weights *W*_struc_ and confidence weights *W*_conf_. These are combined using a user chosen mixing parameter *α*, though we recommend *α* = 0.5, using the following formula:

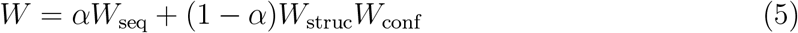

To compute *W*_seq_, we compute the midpoints of all peptides: mid_*i*_. The sequence distance between two peptides is defined as the absolute distance between their midpoints: *d*_*ij*_ = |mid_*i*_ − mid_*j*_|. If the overlap between two peptides *i* and *j* is non-zero we compute their overlap fraction as

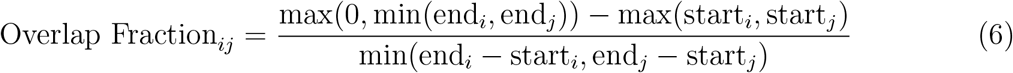

where end_*i*_, start_*i*_ are the end and start of the peptide respectively. The overlap fraction quantifies how much two peptides share common amino acid residues. A value of 1 indicates complete overlap of the shorter peptide with the longer one, while 0 indicates no shared residues. Peptides with higher overlap fractions with have more correlated measurements since they probe the same protein regions. The sequence weights are then given by:

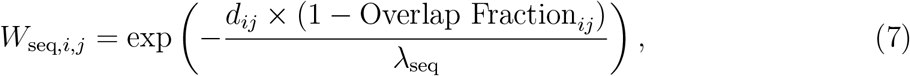

where *d*_*ij*_ is the distance between peptide midpoints and *λ*_seq_ is a scaling parameter controlling the decay of rate of sequence-based correlations (we recommend 1 in practice). The structure weights *W*_struc_ are computed by first obtain the centroids for every peptide. This creates stronger correlations between peptides that are both close in sequence and have significant overlap. The term (1 - Overlap Fraction) ensures that overlapping peptides maintain high correlation even if their midpoints are distant, while non-overlapping peptides have correlation that decays exponentially with sequence distance. The structural distance is then the euclidean distance between the centroids *d*_struc,*i,j*_ = ∥centroid_*i*_ − centroid_*j*_∥_2_. The structural weights are then given by

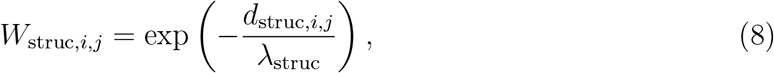

where *λ*_struc_ is user chosen structural decay parameter and we recommend 1 in practice. Peptides that are close together in 3D space will have similar HDX behaviour due to shared local structural environment and solvent accessibility, even if they are distant in the primary sequence. The exponential decay ensures that very distant peptides have minimal structural correlation. For confidence weights, we use the mean pLDDT for each peptide, 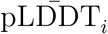 The confidence weights are then given by:

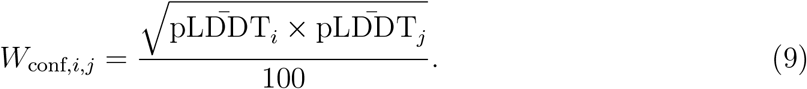

This down-weights correlations involving peptides in poorly predicted structural regions. When both peptides have high confidence (pLDDT ≈ 90 − 100), the weight approaches 1. If either peptide is in a low-confidence region, the structural correlation is reduced accordingly, making the method robust to prediction errors. The resultant weight matrix *W* is normalised row-wise

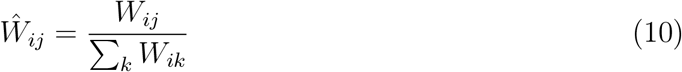

This ensures each peptide’s total correlation weight sums to 1, preventing peptides in dense regions from dominating the correlation structure. Each row represents how a peptide’s correlation is distributed among all other peptides. The correlation matrix is then obtained by

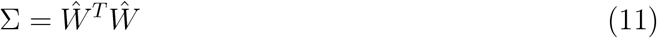

The matrix multiplication transforms direct peptide-to-peptide weights into a proper correlation matrix. We then perform diagonal normalisation by computing the square root of the diagonal entries of this matrix (*d*_1_, …, *d*_*N*_) and dividing each entry Σ_*ij*_ by *d*_*i*_*d*_*j*_. We finally ensure the matrix is symmetric by summing it with it’s transpose and dividing by 2. This ensures peptides perfectly correlate with themselves and removes any numerical asymmetries that arise from computation.

## Code availability

The package is available of GitHub: https://github.com/ococrook/hdx-sFDR.

## Supporting information

Supplementary manuscript

## Acknowledgments

OMC acknowledges funding from a Todd-Bird Junior Research Fellowship and MRC Fellowship MR/Y010078/1.

## Notes

### Competing Interest Statement

OMC is on the scientific adivisory board of Evolvere Bioscience Ltd. He has recevied funding from Pelago Biosiences and GlaxoSmithKline. These edities had no involvement in this work.

