## Supplementary manuscript for "Structure guided significance testing correction for hydrogen deuterium exchange mass spectrometry"

Oliver M. Crook *[1]

[1]Kavli Institute for Nanoscience Discovery, Department of Chemistry, University of Oxford, Oxford, UK

June 16, 2025

### 1 Full epitope mapping examples

*

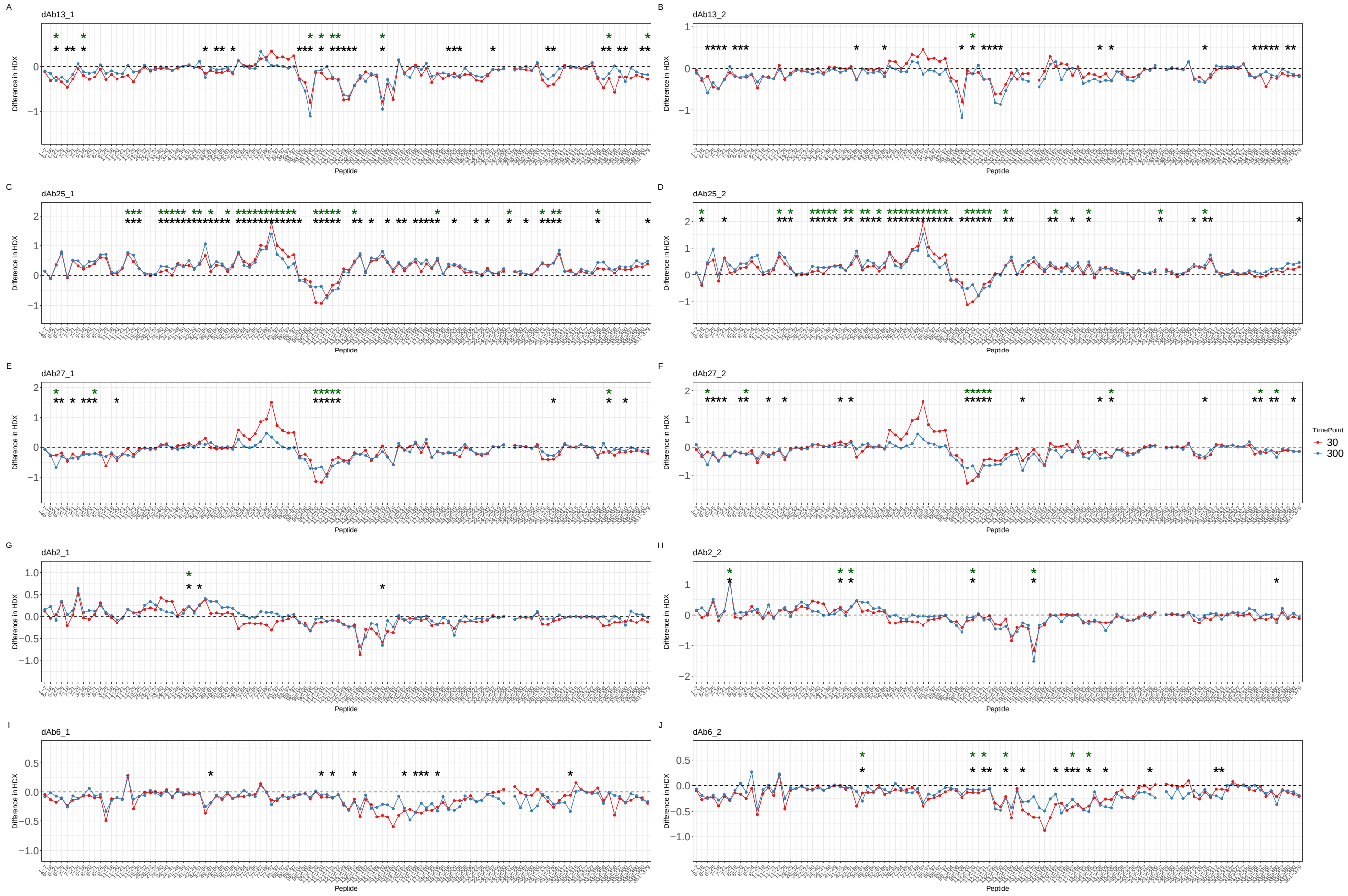

Figure 1: **HOIP Epitope mapping experiments** T butterfly plot with peptides mapped against pointwise deuterium uptake differences in units of Daltons. Structural regions are highlighted using a colour bar. The dark green stars indicate peptides that are significant using standard FDR procedures whilst the black stars indicate peptides that are significant using the sFDR approach. The difference in HDX is between unbound HOIP-RBR and HOIP bound to various dAbs.
